# sRQA: An Integrative Pipeline For Symbolic Recurrence Quantification Analysis

**DOI:** 10.64898/2026.03.31.715624

**Authors:** Austen Curtin, Emma Merriman, Paul C.P. Curtin

## Abstract

Recurrence Quantification Analysis (RQA) is a powerful phenomenological method for characterizing dynamical systems from sequential empirical data, but it is fundamentally limited to continuous signals. Symbolic RQA (sRQA) extends this framework to discrete state sequences, enabling the analysis of both inherently discrete systems and continuous systems where state-based dynamics and motifs are of interest. Despite its promise, accessible and unified software support for sRQA has remained limited. Here we introduce the sRQA package, an open-source R library that consolidates discretization and symbolization, data visualization, and computation of recurrence and cross-recurrence metrics into a single accessible toolset. We validated the method using simulated data with known dynamical properties, confirming that sRQA metrics behaved as theoretically expected. We then demonstrated the utility of sRQA across three real-world applications. First, we applied sRQA to ECG recordings, showing that symbolic recurrence metrics reliably distinguished atrial fibrillation from normal sinus rhythm, with an XGBoost classifier achieving 92% accuracy and an AUC of 0.97. Second, we applied sRQA to fMRI BOLD time series from the dorsal attention network, finding that symbolic and cross-recurrence metrics differentiated movie-viewing from resting-state conditions, revealing greater regularity and inter-subnetwork coordination during task engagement. Third, we applied sRQA to intrinsically symbolized sequences of pauses in speech, identifying valence-specific differences in pause dynamics between truthful and deceptive statements, as well as sex differences in pause structure during negatively-valenced speech. Together, these results demonstrate that sRQA provides a flexible and sensitive framework for characterizing discrete and discretized dynamical systems across biological and behavioral domains.

**AUTHOR SUMMARY:** Many biological and behavioral systems are best understood as sequences of discrete states rather than smooth, continuous processes. For example, a heartbeat that shifts between rhythms, a brain that transitions between activity patterns, or a speaker who pauses and resumes in ways that carry meaning. Standard methods for analyzing the dynamics of such systems were not designed with this kind of data in mind. Here, we introduce the sRQA package, an open-source software library that makes it straightforward to apply symbolic recurrence analysis to both discrete and continuous data. We demonstrate the library across four examples: simulated data with known properties, cardiac recordings distinguishing atrial fibrillation from normal heart rhythm, brain imaging data capturing differences between rest and task engagement, and speech recordings where pause patterns differ between truthful and deceptive statements. In each case, sRQA revealed meaningful structure in the data that would be difficult to detect with conventional tools. We hope this library will make symbolic recurrence analysis more accessible to researchers across the biological and behavioral sciences.

## INTRODUCTION

Temporal dynamics in physiology and behavior are commonly studied through the lens of dynamical systems theory, which provides a robust computational framework for the analysis of temporal structure (1,2). This perspective invites two general methodological approaches to the analysis of biological systems, consisting either of a classical, theoretical approach, or a data-driven, phenomenological approach (1,3). In practice, both perspectives deal with the analysis of continuous time series. This presents an inherent challenge in the analysis of biological sequences that are discrete in nature, particularly those commonly encountered in molecular, physiological, and behavioral data. Here we introduce the *sRQA* library to provide a computational tool for the analysis of temporal dynamics in discrete biological sequences.

Discrete data structures emerge in the analysis of biological data either in consideration of inherently discrete states (3), or when continuous measures may be more easily interpreted or analyzed as discrete states. Examples of inherently discrete data types include sequences of binary events, as are observed in the analysis of action potential “spike trains”; analysis of genetic, epigenetic, proteomic, and/or similar molecular data, consisting of nucleotide sequences; and, aspects of natural language processing involving analysis and quantification of word sequences (4,5). Biological sequences that may be practically treated as discrete sequences include examples such as behavioral sequences, data that are ranked or treated with quantile-based ‘binning’, and/or frequency data treated with spectral analysis for tonal categorization. Discrete data structures thus emerge across multiple physical and temporal scales of biological organization.

Discrete data nonetheless present a fundamental challenge for classical dynamical methods based on the specification of a set of ordinary or partial differential equations (ODE/PDEs) (2) which fundamentally deal with the dynamics of numeric systems. Phenomenological approaches offer an alternative by grounding analysis in the observable behavior of a system rather than in assumptions about its underlying mechanics (6). Rather than specifying ODEs, these methods characterize a system’s behavior through the statistical properties of its time series data, making them essentially descriptive in nature. A key strategy in phenomenological analysis is reconstructing the underlying attractor – the structure in state space toward which the system’s trajectory tends to evolve – from observed data. The theoretical basis for this is Takens’ embedding theorem, which shows that a scalar time series can be used to reconstruct a state space that preserves the essential properties of the original system (7). Complementary approaches, such as potential energy landscape methods, offer related ways of visualizing and quantifying the stability of system states (8). Together, these techniques allow researchers to study the structure of a dynamical system without ever having access to its governing equations.

Recurrence Quantification Analysis (RQA) is a canonical, phenomenological method for studying continuous dynamical systems. RQA operates by first constructing a recurrence plot which is a two-dimensional representation of the times at which a system returns to similar regions of its state space (9). These plots have a characteristic structure: isolated points reflect random or uncorrelated fluctuations, diagonal lines reflect periodic or deterministic dynamics, and vertical and horizontal lines reflect the system dwelling in a particular state. Quantitative measures are then extracted from these patterns, allowing researchers to characterize the attractor structure reconstructed from observed time series data. Metrics such as determinism and average diagonal line length capture the degree of periodicity and predictability in system dynamics, while measures such as laminarity quantify the tendency of a system to become trapped in particular states. RQA has been applied to characterize elemental metabolic dynamics across development, identifying early signatures that distinguish neurodevelopmental conditions such as autism spectrum disorder (ASD) and attention-deficit/hyperactivity disorder (ADHD) from typical development (10–14). This framework has also been extended to network-level analyses of elemental metabolism (15), and to neuroimaging data, where RQA captured altered periodic dynamics in the default mode network in ASD and ADHD, and associations between elemental metabolic dynamics and functional connectivity (16,17). Despite its utility, however, RQA is fundamentally limited to continuous time series data which is a significant constraint given that many biological systems are better described as sequences of discrete states.

Symbolic RQA adapts the RQA framework for sequences of discrete states rather than continuous time series data (18–21). This makes it applicable to systems that are inherently discrete, but also opens up a distinct mode of analysis for continuous systems (22), with the additional benefit that this approach avoids the need for estimating parameters necessary for the implementation of RQA, such as a recurrence threshold, which may be computationally challenging and/or lack theoretical justification. By first discretizing a time series, researchers can study state-based dynamics and motifs, that is, repetitive chunks of sequences that may reflect meaningful behavioral or physiological patterns. Several approaches to discretization exist, including ranking-based and ordinal methods, each of which maps continuous values onto a finite set of symbols according to different criteria (23–25). Ordinal methods in particular have known shortcomings, however; most notably, their sensitivity to window size, which can substantially affect the resulting symbolic sequence and the dynamics it appears to reflect (26–29). Despite the promise of sRQA, to our knowledge support for this approach remains limited in established analytical libraries, creating a practical barrier to its wider adoption (30–33).

To address this gap, we developed the sRQA package, an open-source library that provides a simple, unified framework for symbolic recurrence analysis. The library integrates three core functions: discretization and symbolization of time series data, data visualization, and computation of recurrence and cross-recurrence metrics. By consolidating these steps into a single accessible toolset, sRQA lowers the barrier to applying symbolic recurrence analysis across a wide range of research contexts. Here, we demonstrate the utility of sRQA across several synthetic and real-world examples. First, we validate the method using simulated data with known dynamical properties. Second, we apply sRQA to ECG recordings to distinguish atrial fibrillation from normal sinus rhythm, demonstrating its utility for characterizing cardiac dynamics. Third, we apply sRQA to fMRI data from the dorsal attention network, using symbolic and cross - recurrence metrics to differentiate task-based and resting-state conditions. Finally, we apply sRQA to intrinsically symbolized speech pause-patterns to examine how the dynamics of pausing differ between truthful and deceptive statements across positive and negative valence contexts.

## RESULTS

We present results across four cases that progressively demonstrate sRQA in different contexts: (1) a simulation study establishing the method’s sensitivity to known dynamical regimes, (2) classification of atrial fibrillation from ECG-derived R-R intervals, (3) characterization of task-related BOLD signal dynamics in the dorsal attention network, and (4) analysis of pause patterns in deceptive speech. Each case employs a different symbolization strategy suited to the data type, illustrating the flexibility of the sRQA framework.

### Case 1. Illustrative Example and Simulation

First, we introduce the core sRQA measures through illustrative synthetic examples. We generated three canonical time series: (1) a stochastic process (white noise), (2) a periodic process (sinusoidal signal), and (3) a chaotic-to-periodic regime shift. Each series was symbolized using the *run length encoding* (RLE) method, which was used to discretize data into one of six discrete symbolic pattern types: Increasing, Decreasing, Peak, Valley, Step Up, and Step Down. To achieve this, data are windowed, and within each window first-order differences are calculated for each data point, yielding a sequence of (+1, -1, or 0) values indicating its change (increase, decrease, or stationary) relative to the preceding points. A run is then defined as a sequence of consecutive values, and an arbitrary number of patterns may be specified by combinatorial sequences of runs. A single positive run, for example, reflects a pattern of monotonic increase, while a sequence of negative then positive runs reflects a valley in the data. A majority-rules voting method is used to assign each point a given symbol type when the application of sliding windows yield discordant symbol assignment.

Figure 1 presents the symbolized time series (a) and symbolic recurrence plots (b) generated in this approach; Table 1 reports all sRQA measures (see Recurrence and Cross-Recurrence Measures in METHODS for definitions) derived from the analysis of these matrices.

**Figure 1.**
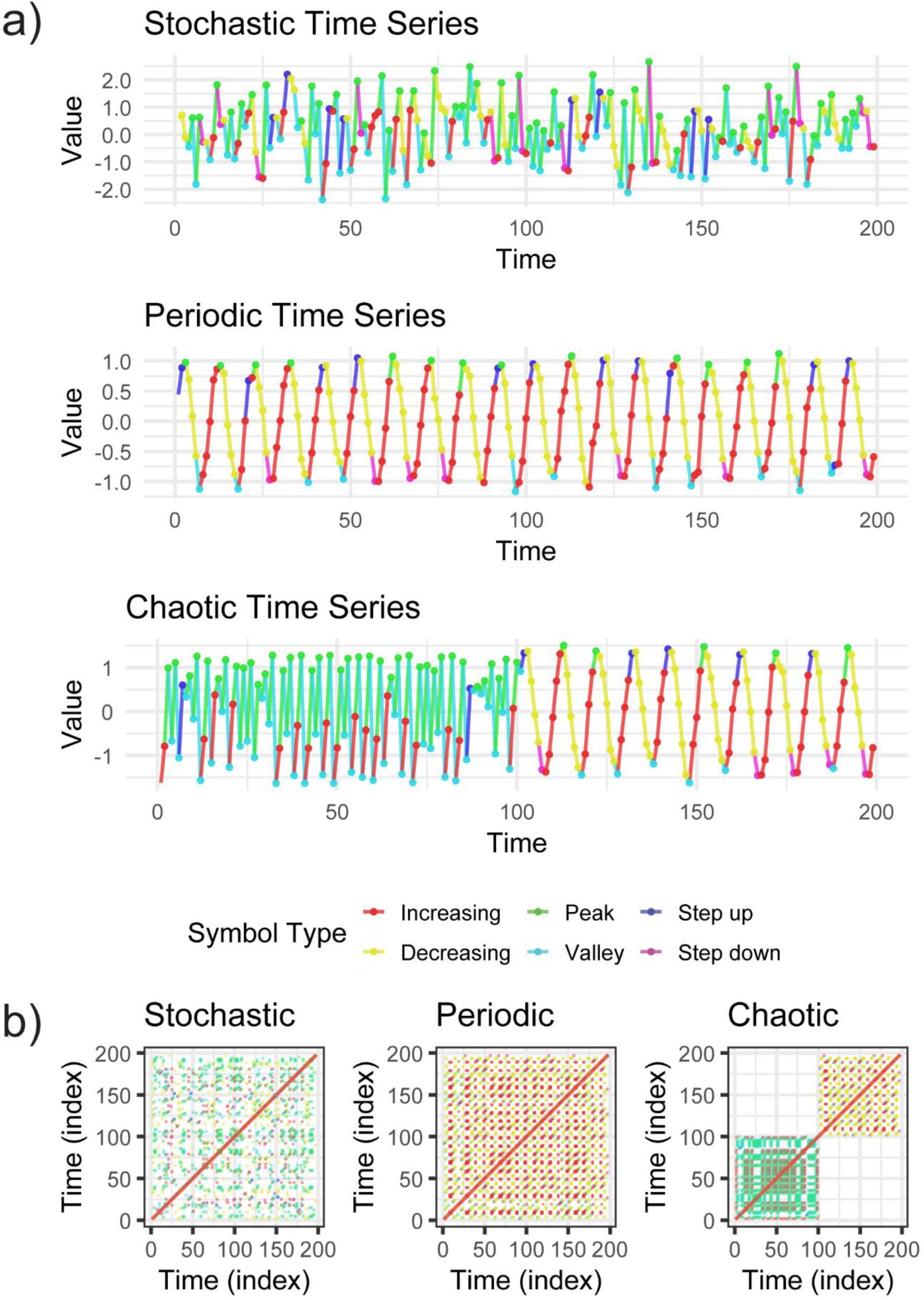
Symbolized time series and symbolic recurrence plots for three canonical dynamical systems. (a) Each time series is color-coded by symbol type: Increasing (red), Decreasing (yellow), Peak (green), Valley (cyan), Step Up (blue), and Step Down (purple). The stochastic system (top) is white noise, the periodic system (middle) is a sinusoidal signal (f = 0.1, σ = 0.1), and the chaotic system (bottom) consists of a logistic map (r = 3.9) for the first 100 observations followed by the sinusoidal signal for the remaining 100. (b) Corresponding symbolic recurrence plots constructed with embedding dimension m = 3. The stochastic system shows sparse, unstructured recurrences; the periodic system displays regular diagonal and grid-like patterning; and the chaotic system exhibits a clear block structure, with dense recurrences within each regime and a visible transition at the series midpoint.

**Table 1.** Symbolic Recurrence Quantification Analysis (sRQA) measures for the three illustrative time series. Measures were computed using *RLE* symbolization.

| Measure | Stochastic | Periodic | Chaotic |
| --- | --- | --- | --- |
| RR | 0.04 | 0.09 | 0.07 |
| DET | 0.67 | 0.66 | 0.86 |
| L | 2.70 | 3.73 | 4.04 |
| Lmax | 7 | 17 | 21 |
| DIV | 0.14 | 0.06 | 0.05 |
| ENTR | 1.15 | 1.48 | 1.81 |
| LAM | 0 | 0.61 | 0.24 |
| TT | 0 | 2.30 | 2.39 |
| Vmax | 0 | 3 | 3 |
| VENTR | — | 0.61 | 0.67 |
| MRT | 18.44 | 14.27 | 7.42 |
| RTE | 3.43 | 1.69 | 2.24 |
| NMPRT | 128 | 981 | 416 |
| TREND | 0.02 | -0.06 | -0.85 |

*Note. RR = recurrence rate; DET = determinism; L = mean diagonal line length; Lmax = maximum diagonal line length; DIV = divergence ( $1/L_{\max}$ ); ENTR = Shannon entropy of diagonal line lengths; LAM = laminarity; TT = trapping time; Vmax = maximum vertical line length; VENTR = Shannon entropy of vertical line lengths; MRT = mean recurrence time; RTE = recurrence time entropy; NMPRT = number of recurrence time points; TREND = trend of recurrence rate away from the main diagonal. An em dash (—) indicates the measure was undefined due to the absence of vertical line structures.*

We found that sRQA measures differentiated the three systems in theoretically expected ways (Table 1). Recurrence rate (RR) was highest for the periodic system (0.09) and lowest for the stochastic system (0.04), while determinism (DET) was highest for the chaotic system (0.86), indicating sustained rule-governed dynamics. Diagonal line measures (L, Lmax, ENTR) increased monotonically, with DIV correspondingly decreasing from stochastic to chaotic, with entropy highest for the chaotic system (1.81), reflecting its mixture of dynamical regimes. Laminarity and related vertical line measures (LAM, TT, Vmax, VENTR) were highest for the periodic system (LAM = 0.61) and lowest for the stochastic system (LAM = 0), clearly differentiating unstructured noise from systems with even transient order. Mean recurrence time (MRT) was longest for the stochastic system (18.44) and shortest for the chaotic system (7.42), with recurrence time entropy (RTE) lowest for the periodic system (1.69), reflecting highly regular return times. Finally, TREND was near zero for the stochastic (0.02) and periodic (−0.06) systems but strongly negative for the chaotic system (−0.85), clearly signaling the regime shift at the series midpoint.

To evaluate the robustness of these patterns, we repeated the analysis across 100 iterations of each system, generating new stochastic, periodic, and chaotic time series on each iteration. Figure 2 presents the distributions of three key sRQA measures, RR, DET, and ENTR, across the three systems (a), windowed sRQA trajectories illustrating regime shift detection (b), and pre- versus post-shift comparisons within the chaotic system (c). Values were standardized for visualization; all statistical tests were conducted on raw measures.

**Figure 2.**
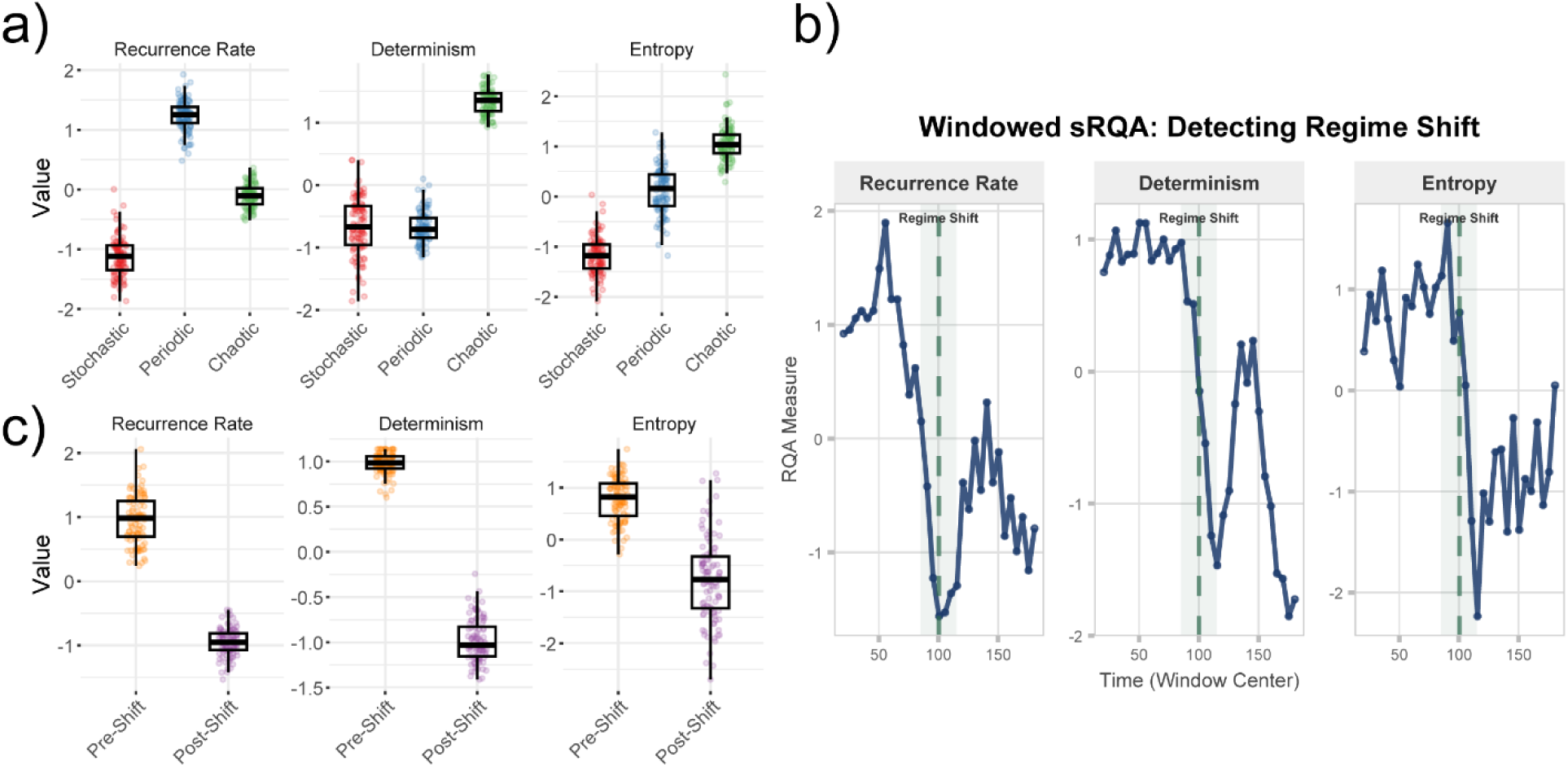
Simulation results across 100 iterations for three sRQA measures: recurrence rate, determinism, and entropy. All values were standardized for visualization. (a) Distributions of each measure across the stochastic (red), periodic (blue), and chaotic (green) systems. All measures differed significantly among systems (Kruskal-Wallis, p < .001), with pairwise comparisons revealing large effect sizes for all contrasts except determinism between the stochastic and periodic systems (d = 0.07, adjusted p = 0.39). (b) Windowed sRQA applied to the chaotic system (window width = 40, step size = 5), with measures plotted as a function of the window center. The dashed green line marks the regime shift at index 100; all three measures show clear transitions near the midpoint. (c) Pre-shift (orange) versus post-shift (purple) distributions within the chaotic system. All measures decreased significantly following the transition from chaotic to periodic dynamics (paired t-tests, p < 0.001), with the largest effect observed for determinism (d = −7.48).

Kruskal-Wallis tests confirmed significant differences among the three systems for all three measures (*p* < .001 in each case). Pairwise comparisons with FDR-corrected Wilcoxon rank-sum tests revealed that all system pairs differed significantly for RR and ENTR (all FDR *Q* < .001), with large effect sizes (*d* ranging from −7.99 to 5.87 for RR and −6.35 to −2.34 for ENTR). For DET, the stochastic and periodic systems did not significantly differ from one another (*Q* = 0.39, *d* = 0.07), but both differed markedly from the chaotic system (*Q* < 0.001, *d* = −5.55 and −8.84, respectively), replicating the single-iteration finding that determinism is especially diagnostic of chaotic dynamics.

Windowed sRQA applied to the chaotic system (b) tracked the regime shift at the series midpoint, with all three measures showing clear transitions near the point of the regime shift at index 100. Formal pre- versus post-shift comparisons confirmed substantial changes in all measures: RR decreased by 45.6% (pre-shift mean = 0.174, post-shift mean = 0.095; paired t-test *p* < 0.001, *d* = −4.37), DET decreased by 30.9% (pre-shift mean = 0.977, post-shift mean = 0.674; paired t-test *p* < 0.001, d = −7.48) and ENTR decreased by 20.3% (pre-shift mean = 1.875, post-shift mean = 1.494; paired t-test *p* < 0.001, *d* = −1.66). Notably, the pre-shift chaotic regime exhibited higher recurrence rate than the post-shift periodic regime, reversing the pattern observed in panel (a). This reflects two factors: the logistic map at *r* = 3.9 visits a concentrated region of the symbol space, producing more repeated symbolic states and thus greater recurrence, and the shorter segment length (100 vs. 200 points) yields fewer opportunities for symbol pattern recurrence in the periodic regime, depressing its RR relative to the full periodic system in panel (a). These findings demonstrate that sRQA reliably differentiates distinct dynamical regimes across repeated iterations and that windowed sRQA can detect spontaneous regime shifts. Overall, these patterns establish the interpretive framework for the application of sRQA in the empirical cases that follow.

### Case 2. ECG

Across all thirteen sRQA metrics, R-R intervals symbolized using the quantile method differed significantly between AF and normal sinus rhythm (*p* < 0.001). AF was associated with significantly lower DET, ENTR, LAM, L, Lmax, Vmax, VENTR, MRT, TT, and NMPRT compared to normal sinus rhythm, which is consistent with the predictable, repeating patterns we would expect to see in a healthy cardiac rhythm. Conversely, RTE, TREND, and DIV were significantly higher in AF, reflecting both the unpredictability and nonstationarity that characterize the condition. Figure 3 (a, b) provides an illustrative example of a symbolized normal sinus rhythm and AF signal respectively, with Figure 3c displaying the standardized beta coefficients for each metric as an ordered differences plot. All measures survived FDR correction.

**Figure 3.**
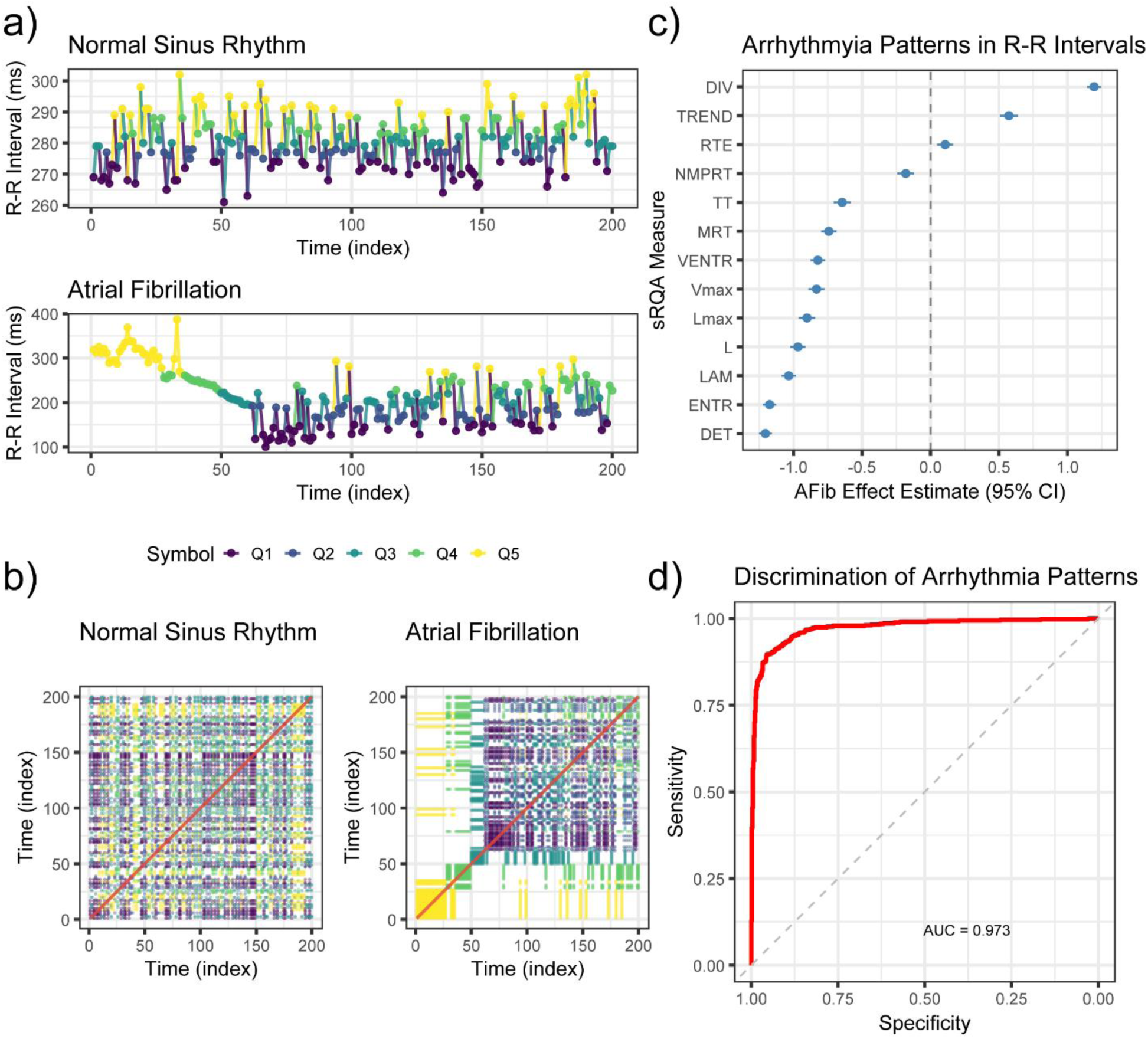
Symbolic recurrence quantification analysis of R-R intervals from normal sinus rhythm and atrial fibrillation (AF) recordings. (a) Example symbolized R-R interval time series for a normal sinus rhythm (top) and AF (bottom) recording, color-coded by quantile symbol (Q1–Q5). The normal sinus rhythm signal displays relatively stable intervals with gradual fluctuations, whereas the AF signal shows irregular, non-stationary variability. (b) Corresponding symbolic recurrence plots. The normal sinus rhythm plot exhibits structured diagonal and vertical patterns indicative of deterministic, repeating dynamics, while the AF plot shows sparser, more fragmented recurrence structures. (c) Standardized beta coefficients (with 95% confidence intervals) from linear mixed effects models comparing AF to normal sinus rhythm across all 13 sRQA measures. Measures below zero (DET, ENTR, LAM, L, Lmax, Vmax, VENTR, MRT, TT, NMPRT) were significantly lower in AF, consistent with reduced predictability; measures above zero (DIV, TREND, RTE) were significantly higher in AF, reflecting greater unpredictability and nonstationarity (all p < .001). (d) Receiver operating characteristic curve for the XGBoost classifier trained on sRQA features to discriminate AF from normal sinus rhythm on the hold-out test set (AUC = 0.973; accuracy = 92.16%).

We found these metrics highly effective for classification of AF relative to normal sinus rhythm. An XGBoost classifier demonstrated excellent performance in distinguishing AF from normal sinus rhythm R-R intervals using sRQA-derived features in a hold-out test set (Figure 3d). The model achieved an overall accuracy of 92.16% (95% CI: 0.90, 0.94), with sensitivity of 90.49%, specificity of 93.49%, positive predictive value of 91.73%, and negative predictive value of 92.48%. The area under the ROC curve was 0.97, indicating excellent discriminative ability.

### Case 3. fMRI

Task-related differences in sRQA metrics were examined across the dorsal attention network (Figure 4a) using linear mixed models, with the BOLD signals symbolized using the quantiles method. In sub-network A (Figure 4b), nine of thirteen metrics differed significantly (*p* < 0.05) between movie-viewing and rest conditions, with all results surviving FDR correction. Movie-viewing was associated with higher LAM, DET, MRT, ENTR, L, TT, and VENTR, but lower DIV and NMPRT, reflecting greater regularity and structured temporal dynamics during movie-viewing. DIV had the most marginal effect (*p* = 0.031, *Q* = 0.045). We found similar results in sub-network B (Figure 4c), with twelve of the thirteen metrics showing significant effects, all surviving FDR correction. Consistent with sub-network A, movie-viewing increased LAM, DET, MRT, RTE, L, ENTR, Lmax, Vmax, TT, and VENTR while decreasing DIV and NMPRT, further supporting the emergence of more predictable, recurrent neural dynamics during movie task engagement. Only TREND was not significant (*p* = 0.273).

**Figure 4.**
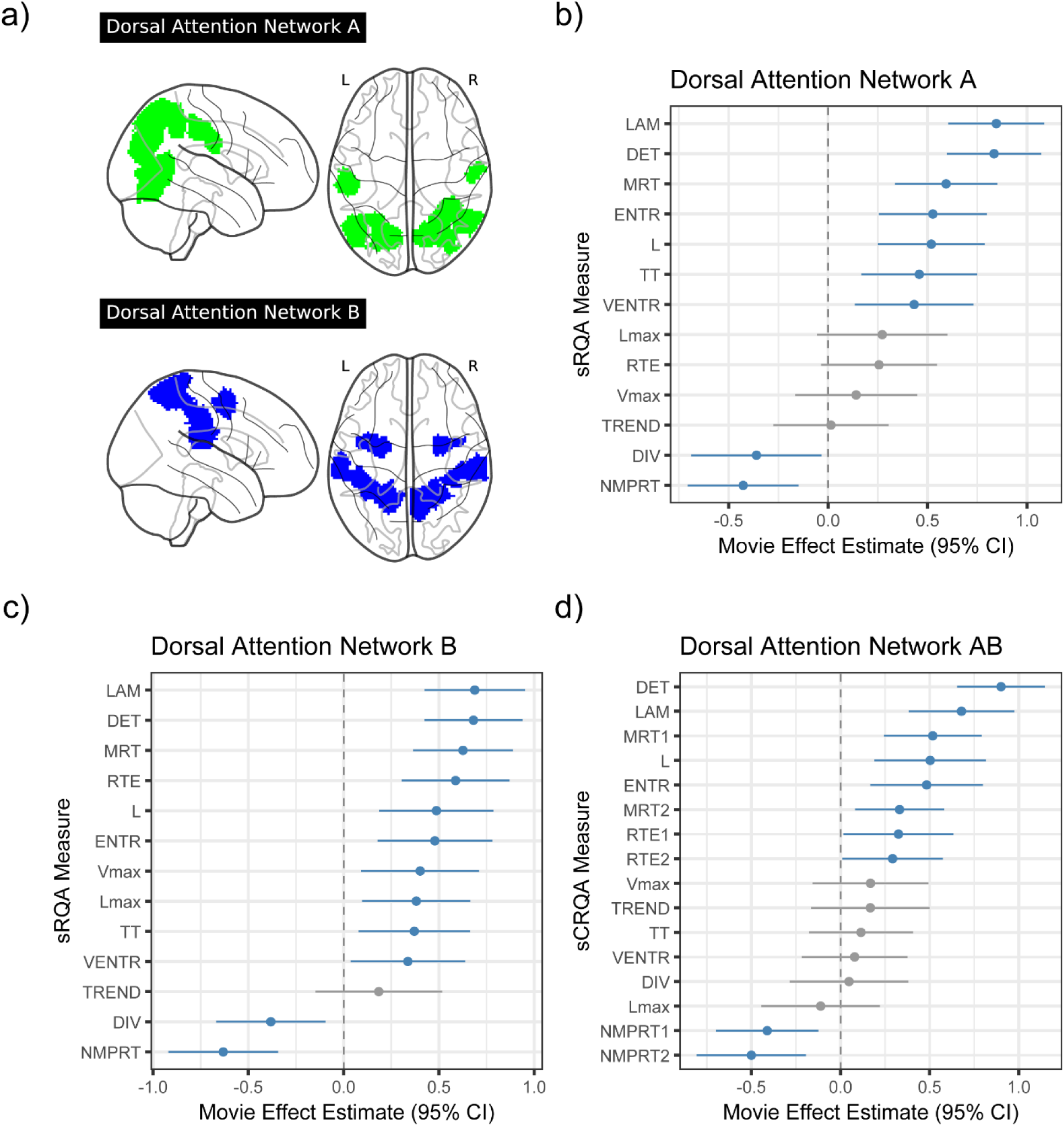
Symbolic recurrence quantification analysis of BOLD signal dynamics in the dorsal attention network during movie-viewing versus rest. (a) Glass brain visualization of dorsal attention sub-networks A (green) and B (blue), displayed in sagittal and axial views. (b) Standardized beta coefficients (with 95% confidence intervals) from linear mixed models comparing movie-viewing to rest for 13 sRQA measures in sub-network A. Blue points indicate significant effects (p < 0.05); gray points indicate nonsignificant effects. Movie-viewing was associated with higher LAM, DET, MRT, ENTR, L, TT, VENTR, and lower DIV and NMPRT, reflecting more structured temporal dynamics during task engagement (all survived FDR correction). (c) Corresponding results for sub-network B, where 12 of 13 measures showed significant task-related differences in the same direction as sub-network A (all survived FDR correction). (d) Symbolic cross-recurrence quantification analysis (sCRQA) between sub-networks A and B. Movie-viewing increased cross-network DET, LAM, MRT1, L, ENTR, and MRT2, while decreasing NMPRT1 and NMPRT2 (RTE1 and RTE2 did not survive FDR correction), indicating stronger inter-network coordination and shared temporal structure during movie-viewing relative to rest.

We also applied cross-recurrence analysis to quantify shared dynamics between symbolic sequences in each network during rest and movie-viewing. Because the cross-recurrence matrix is asymmetric, certain measures are computed separately for each sequence, yielding separate measures for each (e.g., MRT1 and MRT2). Cross-recurrence analysis between sub-networks A and B (Figure 4d) revealed task-related differences in how the two sub-networks coordinated with one another, with eight of sixteen sCRQA metrics showing significant effects that survived FDR correction. Movie-viewing increased cross-network DET, LAM, L, MRT1, MRT2, and ENTR, while decreasing NMPRT1 and NMPRT2, suggesting that the two sub-networks developed stronger coordination and more shared temporal structure during task engagement compared to rest. RTE1 and RTE2 were significant at uncorrected thresholds (*p* < 0.05) but did not survive FDR correction (adjusted *Q* = 0.067 for both). No significant differences emerged for Lmax, DIV, VENTR, TT, Vmax, or TREND.

### Case 4. Speech

As illustrated in Figure 5a, an initial analysis of speech data quantified the interval of pauses between words in participants’ speech; these were then binarized, creating a binary vector corresponding to sequences of intervals with pauses (coded as 1) or no pauses (coded 0) between words. We applied sRQA to this binary sequence of pause sequences (Figure 5a, 5b) in each experimental condition, and tested for interactions between experimental conditions (valence, veracity) and participant characteristics (sex). Four significant valence x veracity interactions emerged across the fourteen sRQA metrics, including ENTR, Vmax, RR, and DIV (Figure 5c); specifically, in all four interactions we found that the effect of truthful speech was specific to positive contexts. In assessing stability across conditions, we found that divergence in some conditions (DIV, β = 0.496, 95% CI: (0.098, 0.893), *p* = 0.015, *Q* = 0.125), such that truths showed greater divergence than lies in positive contexts (*p* = 0.002), but not in negative contexts. The complexity of pause patterns, similarly, tended to differ across experimental conditions (ENTR, β = -0.460, 95% CI: (-0.839, -0.080), *p* = 0.018, *Q* = 0.125), such that lies showed greater entropy than truths in positive contexts (p = 0.004), but not in negative contexts. We also found that the persistence of stable states tended to differ across experimental conditions (Vmax, β = -0.416, 95% CI: (-0.826, - 0.006), *p* = 0.047, *Q* = 0.158), such that lies showed longer sustained vertical structures than truths in positive contexts (*p* = 0.01), but not in negative contexts. Last, we found that overall patterns of repetition tended to differ across experimental conditions (RR, β = -0.389, 95% CI: (-0.774, -0.004), *p* = 0.048, *Q = 0.158*); such that lies showed greater recurrence than truths in positive contexts (p = 0.011), but not in negative contexts. Despite the consistency of these effects, in particular the notable valence-dependent differences that emerged in the truthful statement condition, none of these multiplicative effects survived FDR correction (*Q* > 0.05).

**Figure 5.**
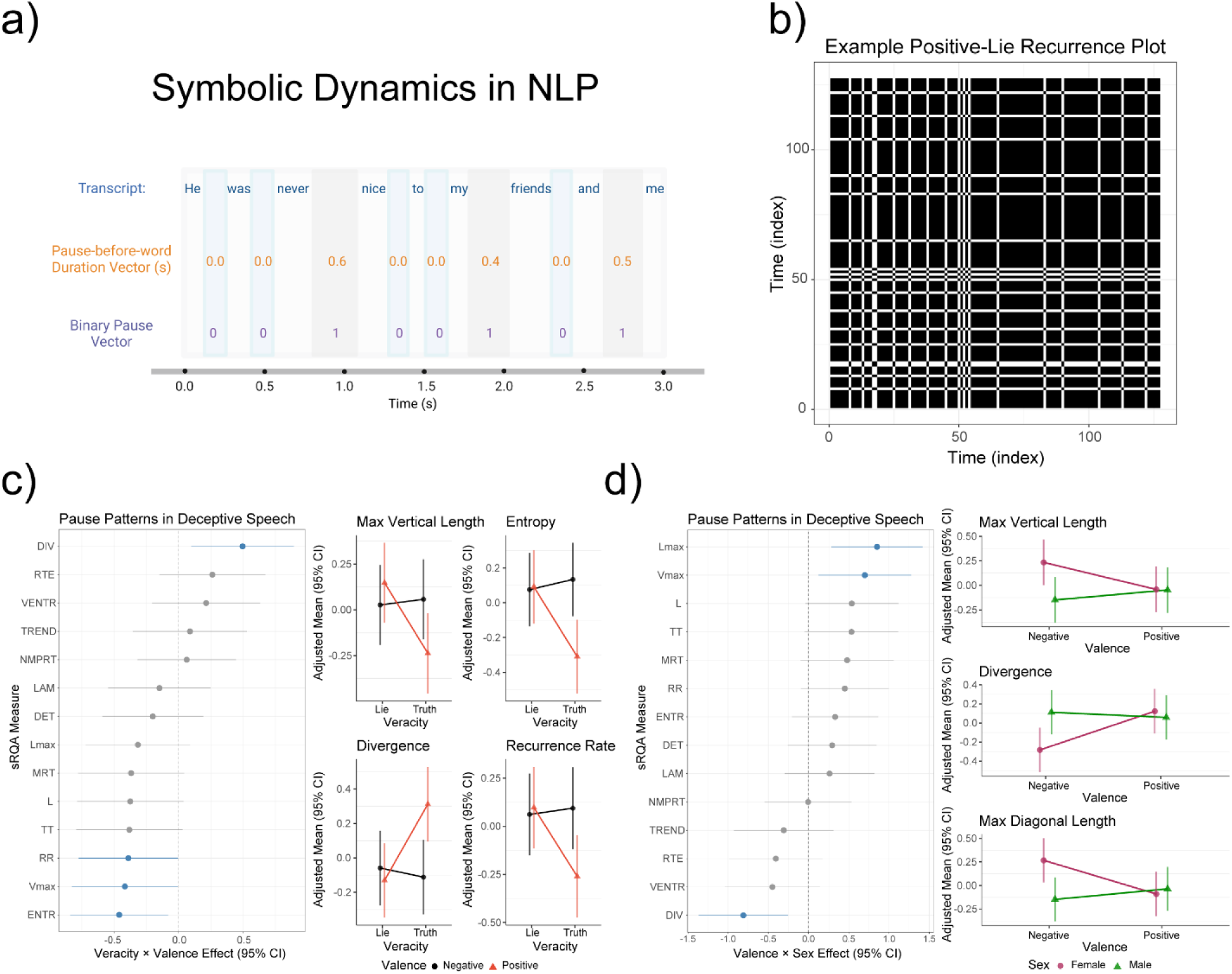
Symbolic recurrence quantification analysis of pause dynamics in deceptive and truthful speech. (a) Illustration of the binarization procedure for speech pause sequences: each word in a transcript is assigned a pause-before-word duration (in seconds), which is then converted to a binary pause vector (1 = pause present, 0 = no pause). (b) Example symbolic recurrence plot from a positive-valence lie trial, constructed from the binarized pause sequence. The dense block structures reflect sustained periods of recurring pause patterns. (c) Veracity × valence interaction effects from linear mixed models across 14 sRQA measures. The left panel displays standardized beta coefficients with 95% confidence intervals (blue = significant at p < 0.05; gray = nonsignificant). The right panels show adjusted means for four significant interactions (Vmax, ENTR, DIV, and RR), illustrating that veracity effects on pause dynamics emerged selectively in positive contexts, with lies showing greater entropy, longer vertical structures, and higher recurrence than truths. (d) Valence × sex interaction effects. The left panel displays standardized beta coefficients as in (c). The right panels show adjusted means for three significant interactions (Vmax, DIV, and Lmax), revealing that in negatively valenced contexts, women exhibited greater pattern stability (higher Lmax and Vmax) and lower divergence than men.

We also examined whether participant sex moderates pause dynamics across valence contexts, finding significant valence x sex interactions involving three metrics: Lmax, Vmax, and DIV (Figure 5d). Much like the veracity findings, all three interactions indicated context-specific effects, but these differed across sexes. Specifically, women exhibited greater sustained periodicity in pause duration than men (Lmax, β = 0.851, 95% CI: (0.287, 1.416), *p* = 0.003, *Q* = 0.031;), but only in conditions of negative valence (*p* = 0.014). We also found that valence-dependent differences in pause stability were sex-dependent and emerged only in negative valence conditions (DIV, β = -0.811, 95% CI: (-1.365, -0.256), *Q* = 0.031), but here divergence was elevated in men relative to women (*p* = 0.019). We found that women tended to exhibit longer sustained non-pause periods (Vmax, β = 0.699, 95% CI: (0.123, 1.275), *p* = 0.018, *Q* = 0.082) in negative valence contexts, but this effect did not survive multiple comparison adjustment.

Finally, we trained an XGBoost classifier to discriminate between truths and lies using sRQA-derived features with leave-one-out cross-validation. The model achieved an AUC of 0.67 and classification accuracy of 0.65, significantly exceeding chance performance (*p* < 0.01), with sensitivity of 66.3%, specificity of 63.8%, positive predictive value of 0.65, and negative predictive value of 0.68.

## DISCUSSION

The four cases presented here demonstrate that sRQA provides a flexible, interpretable framework for characterizing temporal dynamics across diverse data types and domains. We discuss each case in turn, comparing the findings within existing literature and highlighting the unique contributions of the symbolic recurrence approach. In particular, we demonstrate the utility of this approach for distinguishing systemic dynamics and corresponding regime shifts; and, demonstrate the utility of these approaches for distinguishing biologically-meaningful dynamics in physiological systems operating across multiple systems and timescales.

### Case 1. Illustrative Example and Simulation

The simulation study confirmed that sRQA produces theoretically consistent and statistically robust distinctions among canonical dynamical regimes. Recurrence rate and entropy reliably separated all three systems, while determinism specifically distinguished chaotic dynamics from both stochastic and periodic processes, consistent with classical RQA findings on the logistic equation (34) and the broader principle that diagonal line structures reflect deterministic rule-governance rather than mere repetition (35,36). The similar determinism values for the stochastic and periodic systems, despite their vastly different recurrence rates, underscore that these measures capture complementary aspects of dynamical structure, a property that has been emphasized in traditional RQA (3) and is preserved in this symbolic extension.

We further found that windowed sRQA was an effective tool in detecting regime shift in a series with chaotic/periodic dynamics, with all three measures highlighted showing clear transitions and large effect sizes discriminating regimes. The largest effect was observed for determinism (*d* = −7.48), reflecting the fundamental change from chaotic to periodic governing dynamics, while the comparatively smaller effect for entropy (*d* = −1.66) is consistent with both regimes producing moderately complex diagonal line distributions. These results parallel findings from windowed RQA in continuous systems, where time-dependent recurrence measures have been used to detect chaos -order transitions (34,36) and extend them to the symbolic domain. Together, these findings validate sRQA as a tool for characterizing dynamical structure and detecting regime shifts in controlled settings, establishing the foundation for the empirical applications that follow.

### Case 2. ECG

The sRQA metrics derived from R-R intervals reliably distinguished AF from normal sinus rhythm, with the observed pattern of effects highly consistent with known cardiac physiology. AF is characterized by irregular, unpredictable ventricular response, and accordingly we observed significantly reduced determinism, entropy, laminarity, and related metrics in AF, alongside elevated divergence and trend. Collectively these results reflect the loss of structured, recurrent dynamics that characterize normal cardiac rhythm. Prior work that applied sRQA to the same dataset (Physionet MIT-BIH AF) also demonstrated strong classification performance using a logistic regression model, achieving 97.7% accuracy, 97.9% sensitivity, and 97.6% specificity (31). Our XGBoost classifier achieved 92.16% accuracy and an AUC of 0.973, which is broadly consistent with this benchmark. The modest difference in accuracy likely reflects methodological differences rather than any fundamental limitation of the approach: the prior model employed a minimal feature set specifically optimized for a logistic classifier, whereas our implementation deliberately leveraged the full feature set generated by the sRQA implementation in order to fully characterize the potential contribution of each feature, rather than optimize classification performance. Further, the approach to discretizing data used in our application was, for demonstration purposes, based on a quantile approach, which may potentially have been less effective than the ordinal method employed previously. Recurrence-based methods more broadly have shown competitive performance in AF classification, including approaches combining recurrence plots with deep residual convolutional networks (37) and symbolic co-occurrence methods applied to RR intervals (38). Taken together, our results are within the range of established recurrence-based methods, and confirm that the sRQA library produces valid, competitive outputs while offering a substantially more accessible and generalized analytical framework than prior implementations.

### Case 3. fMRI

The sRQA metrics derived from BOLD time series in the dorsal attention network reliably differentiated movie-viewing from resting-state. Across both subnetworks, movie-viewing was associated with greater determinism, laminarity, entropy, and related metrics, reflecting more structured, recurrent, and predictable neural dynamics during task engagement relative to rest. This is broadly consistent with prior work showing that naturalistic stimuli reorganize brain state dynamics away from the diffuse, bistable patterns characteristic of rest toward more structured and temporally coherent sequences of activity (39). The specific involvement of the dorsal attention network in this reorganization aligns with evidence that movie-watching elicits significant changes in DAN connectivity relative to rest (40,41), and that the DAN shows distinct dynamics during naturalistic engagement compared to unconstrained rest (42). The sCRQA results extend these findings by showing that inter-subnetwork coordination was also significantly enhanced during movie-viewing, with greater shared determinism, laminarity, and entropy between subnetworks during task engagement compared to rest. Overall, these results demonstrate that sRQA is sensitive to meaningful task-related differences in neural dynamics, and that sCRQA offers a complementary lens for characterizing inter-network coordination that goes beyond conventional linear connectivity approaches.

### Case 4. Speech

Our application of sRQA to characterize speech dynamics across multiple experimental conditions emphasized the sensitivity of this approach in detecting the influence of psychological dynamics on subtle aspects of speech, particularly pause behavior (43). Though pause characteristics as a marker of deception are a popular metric in speech deception detection, prior studies focused primarily on the frequency, duration, or general summary statistics of pauses, and report inconsistent results. In particular, prior studies report contradictory findings in leveraging pause duration in deception-detection; at least two studies report increased pausing during deception (44,45), while another has reported decreased pausing during deception (46), and another showed no association (47). In contrast to these approaches, which focus on the duration of pause, our analysis focused on the temporal organization of pause patterns, capturing metrics such as recurrence, complexity, and stability over time. Additionally, most deception research examines veracity in negative or criminal content, leaving positive valence contexts largely unexplored (48). The present analysis addresses both gaps, revealing that deceptive and truthful speech differ in the temporal dynamics of their pause patterns, but these effects are critically dependent on higher order interactions among psychological conditions (valence and veracity) as well as sex-dependent effects specifically within negatively-valenced contexts. This is consistent with prior findings that sex moderates vocal characteristics during deceptive speech (49). The complexity of these effects, and the potential involvement of higher order interactions among underlying biological and psychological factors, emphasize the need for future studies with larger samples to untangle these effects with appropriate statistical power. Our findings suggest that sRQA could be a useful tool in this application.

In summary, by organizing symbolization, visualization, and computation of recurrence and cross-recurrence metrics into a single accessible toolset, the sRQA package lowers the practical barrier to adopting symbolic recurrence analysis. The four cases presented here confirm that the method produces theoretically consistent and empirically meaningful results across diverse data types and symbolization strategies. Several directions for future development are worth noting. First, while the present work focused on three symbolization strategies (RLE, quantiles, and intrinsic binarization), the package also supports ordinal methods where smaller window sizes may be appropriate. Second, the combination of sRQA features with machine learning classifiers showed promising but variable performance across cases, emphasizing that this approach can be useful in the application of traditional (associative) statistical approaches as well as predictive pipelines that leverage machine learning methods. Finally, the symbolic recurrence framework is well positioned for application to emerging data types in digital health and behavioral science, including wearable sensor streams, ecological momentary assessment data, and natural language sequences, where discrete or discretized representations are natural and traditional RQA cannot be directly applied. We hope that by making sRQA accessible and well-documented, this package will encourage broader adoption of symbolic recurrence methods and open new avenues for studying the dynamics of complex systems.

## MATERIALS AND METHODS

### Recurrence and Cross-Recurrence Measures

Symbolic recurrence quantification analysis (sRQA) extracts a set of measures from the symbolic recurrence matrix that characterize different aspects of the temporal structure of a discrete state sequence. We briefly define each measure, below, based on the nomenclature and algorithms introduced and/or reviewed in Marwan et al. (2007) (9) and Webber and Zbilut (2005) (3); these metrics are also discussed and implemented for the analysis of continuous time series in the Cross Recurrence Toolbox software implemented in Matlab (50), and the Dynamical Systems Library implemented in Julia (51).

The most basic measure, recurrence rate (RR), is the proportion of recurrent points in the matrix and reflects how frequently the system revisits previously observed symbolic states. Measures derived from diagonal line structures capture the degree of deterministic, rule-governed behavior in the system. Determinism (DET) is the proportion of recurrent points that fall on diagonal lines of at least length *l*min, indicating predictable sequences of recurring states. The mean diagonal line length (L) reflects the average duration of these deterministic episodes, while the maximum diagonal line length (Lmax) captures the longest uninterrupted stretch of predictable behavior. Divergence (DIV = 1/Lmax) provides the inverse perspective, indexing how quickly the system diverges from predictable trajectories. The Shannon entropy of the distribution of diagonal line lengths (ENTR) quantifies the complexity of the system’s deterministic structure, with higher values indicating a more heterogeneous mixture of short and long predictable sequences.

Measures derived from vertical line structures capture the tendency of a system to persist in, or become trapped in, particular states. Laminarity (LAM) is the proportion of recurrent points forming vertical lines of at least length *l*min, and trapping time (TT) is the average length of these vertical structures, reflecting how long the system remains in a given state. The maximum vertical line length (Vmax) and the Shannon entropy of vertical line lengths (VENTR) further characterize the extent and complexity of these laminar episodes.

Recurrence time measures characterize the temporal spacing between successive returns to a given symbolic state. Mean recurrence time (MRT) is the average interval between returns, with longer values indicating less frequent revisitation. Recurrence time entropy (RTE) quantifies the regularity of these return intervals, with lower values indicating more periodic return patterns. The number of recurrence time points (NMPRT) reflects the total number of observed return events. Finally, trend (TREND) measures the drift in recurrence density away from the main diagonal, capturing nonstationarity in the system’s dynamics; values near zero indicate stationarity, while negative values indicate a progressive loss of recurrence structure over time.

For cross-recurrence quantification analysis (sCRQA), the same measures are computed from a cross-recurrence matrix constructed between two symbolic sequences rather than within a single sequence. Because the cross-recurrence matrix is asymmetric, certain measures are computed separately for each sequence. Specifically, mean recurrence time, recurrence time entropy, and number of recurrence time points are computed from both the rows and columns of the cross-recurrence matrix, yielding paired measures indexed by sequence (e.g., MRT1 and MRT2, RTE1 and RTE2, NMPRT1 and NMPRT2) that reflect how frequently each sequence revisits states also visited by the other. The remaining cross-recurrence measures (DET, LAM, L, Lmax, DIV, ENTR, Vmax, VENTR, TREND) are computed from the full cross-recurrence matrix and are interpreted analogously to their single-sequence counterparts, but reflect shared or coordinated dynamics between the two systems rather than within a single system.

### Data Acquisition and Preprocessing

#### Case 1. Illustrative Example and Simulation

Three canonical time series of length *N* = 200 were generated to represent distinct dynamical regimes. The stochastic system consisted of independent draws from a standard normal distribution (μ = 0, σ = 1). The periodic system was a sinusoidal signal with frequency *f* = 0.1 (10 time units per cycle) and additive Gaussian noise (σ = 0.1). The chaotic system comprised two concatenated segments of 100 observations each: the first was generated from a logistic map at *r* = 3.9 (a regime exhibiting fully developed chaos) and the second from the same sinusoidal signal used in the periodic system, producing a regime shift at the series midpoint. Both segments of the chaotic system were independently z-scored prior to concatenation. For the single-iteration analysis, a fixed seed was used for visualization purposes and clarity; for the simulation study (100 iterations), the logistic map initial condition was drawn uniformly from [0.01, 0.99] on each iteration, and new noise samples were generated for the stochastic and periodic systems to capture variability across realizations.

#### Case 2. ECG

We obtained electrocardiogram (ECG) recordings from the PhysioNet MIT-BIH Atrial Fibrillation Database (52,53), a comprehensive publicly available resource for cardiac arrhythmia research. This database comprises recordings from 25 patients, encompassing 149.06 hours of normal sinus rhythm (N), 93.77 hours of atrial fibrillation (AF), and 6.6 hours of other arrhythmias including atrial flutter and atrioventricular junctional rhythm. Each patient’s ECG record was segmented into consecutive, non-overlapping windows of 200 R-R intervals (31). Window classification followed a majority-vote approach: windows with more than 50% of cardiac cycles annotated as AF were labeled as AF, while those with predominantly normal sinus rhythm were labeled as N. Windows dominated by other arrhythmias were designated as O and excluded from the primary analysis.

#### Case 3. fMRI

We obtained preprocessed BOLD time series data from the Healthy Brain Network dataset via the Reproducible Brain Chart repository (54,55). The data were preprocessed using the Configurable Pipeline for Analysis of Connectomes (56) and parcellated into 200 brain regions using the Schaefer atlas, with regions labeled according to the Yeo 17-network parcellation (57,58) and 36 movement and noise artifacts removed. We included only participants with complete, high-quality data as determined by the repository’s quality control standards. Specifically, we queried participants who had both 5 minutes of resting state and movie-watching (Despicable Me) scans in addition to having no clinical diagnosis. This resulted in a total of 71 participants and 142 scans (2 scans per person). We focused on the Dorsal Attention Network (DAN) because it shows distinct connectivity patterns during movie watching compared to rest (40–42,59). Using this parcellation, we identified 22 regions forming the DAN, which were divided into two subnetworks (A and B) as defined in Nilearn v0.12.0 (Python v3.9) (60). We created a single time series for each subnetwork by averaging activity across all regions within that subnetwork.

#### Case 4. Speech

The data for this study comes from the Miami University Deception Detection (MU3D) dataset, a publicly available collection of video recorded speech from 80 individuals (40 women, 40 men, 40 black, 40 white) under laboratory-controlled conditions (61). Participants were instructed to provide both truthful and deceptive descriptions about interpersonal experiences with varied valence. Thus, each participant generated four speech recordings following a 2x2 within-subjects experimental design: statement veracity (lie or truth) x statement valence (positive or negative). So, the final analytic cohort yields 320 observations, with individual videos spanning from 24.33 to 43.6 seconds in length, and word count ranging from 46 to 160 words.

Audio files were extracted from the 320 video recordings using the MoviePy v2.2.1 (Python v3.9) (62). Resulting audio files were transcribed using Assembly AI’s automatic speech recognition (ASR) model, which provides word-level parsed timestamps. Transcription accuracy was evaluated by calculating the Jaccard similarity between Assembly AI and manual transcripts (63). Token level accuracy was an average of 83.1% across 320 transcripts. For each parsed word in the transcripts, the pause duration preceding that token was calculated as the difference in time between the current token’s start time and the previous token’s end time in milliseconds; words with zero difference are considered continuous/fluent speech. For each transcript, a vector of pause duration was generated that aligns with each parsed word sequence. These vectors are sparse, as most participant’s speech is continuous, so pause duration vectors were binarized into a categorical sequence which represented the presence (coded as 1) or absence (coded as 0) of pauses. This binarization produced an intrinsic symbolization appropriate for recurrence quantification analysis.

### Symbolization Methods

The sRQA library implements three approaches for discretizing continuous data, and a fourth approach for inherently discrete data.

#### Rank-Based Symbolization

The simplest method of discretizing a time series is implemented by quantile-based ranking of each point in the time series. The sRQA library allows the specification of any arbitrary quantile scheme – for example, quartiles, quintiles, deciles, etc. – which will yield a corresponding distribution of symbols. This approach may be particularly appropriate in dealing with noisy data, where the absolute value of a given point in the series is less important than its relative magnitude.

#### Ordinal-Based Symbolization

Ordinal pattern symbolization converts a numeric time series into a discrete symbolic sequence by encoding the rank-order structure within sliding windows of length *w*. For each window, the *w* data points are replaced by their rank permutation, and this permutation is mapped to a unique integer symbol. With a window of size *w*, there are *w*! possible ordinal patterns, each capturing a distinct local ordering of values (e.g., for *w* = 3, the six patterns correspond to all possible monotonic, peak, and valley configurations). Ties are broken randomly. Each symbol is assigned to the terminal position of its corresponding window, yielding a symbolic series of length *n* − *w* + 1, where n is the original series length. This approach, rooted in the permutation entropy framework of Bandt and Pompe (2002), is robust to monotonic transformations of the data and captures local dynamic structure without requiring distributional assumptions or parameter-dependent thresholds. A similar approach has been implemented previously in the Cross Recurrence Plot Toolbox (50) framework in Matlab, and this method has also been implemented in a number of empirical analyses (30,32,33,64).

#### Run Length Encoding Symbolization

The run length encoding (RLE) symbolization method introduced here is designed to solve the problem of exploding symbol complexity implicit to the ordinal method; to our knowledge, this is a novel symbolization method that has not previously been employed in symbolic recurrence quantification analysis. More specifically, as noted above, because the number of symbols derived in the ordinal method will be a factorial function of window size, in practice larger window sizes than *w* = 3 or *w* = 4 become uninterpretable. To avoid this, the run length encoding (RLE) symbolization method converts a numeric time series into a discrete symbolic sequence by classifying the directional structure within overlapping sliding windows of length *w*. For each window, first differences are computed and their signs are quantized into positive, negative, or zero (flat) based on an adaptive threshold proportional to the window’s range and a user-specified sensitivity parameter. Run length encoding (65), as commonly applied as an image compression algorithm, is then applied to this sequence of directional signs, yielding a compact representation of the number, order, and direction of monotonic segments within the window. The resulting run structure is mapped to one of up to eight symbols: monotonic increase, monotonic decrease, single peak, single valley, step up, step down, upward oscillation, and downward oscillation. Because each data point falls within multiple overlapping windows, a majority -vote scheme is used to determine the final symbol assignment at each time step, ensuring that every point in the original series receives a symbol. This approach captures local morphological features of the time series without requiring distributional assumptions, and the voting mechanism provides robustness against boundary effects and transient noise.

#### Inherent symbolization

Last, the *sRQA* library readily accepts data that are inherently discrete in nature and therefore require no discretization step.

### sRQA Implementation

#### Case 1. Illustrative Example and Simulation

Each example time series was symbolized using the RLE method with six symbol types (Increasing, Decreasing, Peak, Valley, Step Up, and Step Down) and a sliding window of size *w* = 3. The goal in this approach is to create a symbolization process that functions similarly to ordinal encoding, but that could be extended to windows of arbitrary size without exploding the diversity of potential symbol types. To achieve this, in each window first differences are computed (i.e., the difference from point 1 to point 2) and converted to direction signs (+1,-1, or 0). Run length encoding (25,65) is then applied to that sequence, grouping consecutive identical directions into “runs”. The number and their values determine the shape classification – for example, a single positive run is classified as a “monotonic increasing”, two runs (positive then negative) are classified as a “single peak”, two runs (negative then positive) are classified as a valley, etc.

Symbolic recurrence matrices were constructed with an embedding dimension of *m* = 3, and all recurrence quantification measures were computed with a minimum diagonal and vertical line length of *l*min = 2. For the single-iteration example, all 14 sRQA measures were examined to illustrate the full range of information captured by the method (Table 1). For the simulation study, three commonly reported measures that include RR, DET, and ENTR, were retained for statistical comparison. For the windowed sRQA analysis, a window width of 40 observations and a step size of 5 were used, with the number of symbols increased to 8 across iterations. All sRQA metrics were z-scored prior to visualization.

#### Case 2. ECG

R-R intervals constructed from ECG recordings were symbolized with the *quantiles* method. We divided the time series into five equally probable categories based on the distribution of values (num_symbols = 5). We then constructed symbolic recurrence matrices with an embedding dimension (*m*) and time delay (*d*) of 1 to capture recurrence based on each symbol type. We then quantified 14 distinct features. Recurrence rate (RR) was excluded from further statistical analyses because this value is fixed when using quantile symbolization. This left us with 13 symbolic recurrence features. All sRQA metrics were z-scored prior to modeling and visualization.

#### Case 3. fMRI

Average BOLD time series for DAN subnetworks were symbolized like *Case 2* with the *quantiles* method (num_symbols = 5, *m* = 1, and *d* = 1). From individual subnetwork symbolic matrices, we quantified 14 distinct features. For cross recurrence symbolization, we quantified 6 additional features resulting from the asymmetry in cross-recurrence matrices. After excluding recurrence rate metrics, we were left with 13 features for individual subnetworks and 16 features for their cross-recurrence. All metrics were z-scored prior to modeling and visualization.

#### Case 4. Speech

Binarized pause vectors are intrinsically symbolized, so no further symbolization methods were necessary. For each of the participant’s four pause-based vectors, a symbolic recurrence matrix was constructed with a *m* and *d* of 1 to preserve each parsed word as an individual observation. Recurrence quantification measures were then extracted from these vectors, computing 14 metrics that quantify different aspects of the pause pattern dynamics. All sRQA metrics were z-scored prior to modeling and visualization.

### Statistical Analysis and Machine Learning Classification

#### Case 1. Illustrative Example and Simulation

For the single-iteration example, a descriptive table of all 14 sRQA measures was constructed to characterize each system (Table 1). Differences among the three systems across 100 iterations were tested using Kruskal-Wallis tests, followed by pairwise Wilcoxon rank-sum tests with FDR correction. Effect sizes were quantified using Cohen’s *d* with pooled standard deviations. To evaluate the regime shift within the chaotic system, pre-shift and post-shift segments were symbolized and analyzed independently on each iteration. Pre-post differences were tested using paired-samples t-tests, with effect sizes computed as the mean paired difference divided by the standard deviation of the paired differences. All measures were z-scored within each variable for visualization.

#### Case 2. *ECG*

Data were analyzed using linear mixed-effects models implemented in R (lme4 package). We examined recurrence quantification analysis (RQA) metrics derived from ECG recordings. Cardiac rhythm type (atrial fibrillation vs. normal sinus rhythm) was dummy-coded and entered as a fixed effect, with participant included as a random intercept to account for repeated measures. Separate models were fit for 13 RQA metrics (DET, L, Lmax, DIV, ENTR, LAM, TT, Vmax, VENTR, MRT, RTE, NMPRT, TREND). Models were estimated using restricted maximum likelihood (REML), and 95% confidence intervals were computed for fixed effects using the broom.mixed package. All p-values were adjusted for multiple comparisons using FDR correction.

A machine learning classification model was developed to distinguish atrial fibrillation from normal sinus rhythm using RQA metrics derived from ECG data. Data were split into training (80%) and test (20%) sets. An XGBoost gradient boosting classifier was implemented using the mlr3 framework in R. Hyperparameter optimization was performed using Hyperband tuning with 5-fold cross-validation on the training set, optimizing area under the ROC curve (AUC). The search space included: number of rounds (100–800), learning rate (0.1-0.3), maximum tree depth (3–12), gamma (0–15), and L1/L2 regularization parameters (0-15 each). The final model was trained on the complete training set using optimized hyperparameters and evaluated on the held-out test set. Model performance was assessed using AUC, sensitivity, specificity, and overall classification accuracy via confusion matrix analysis.

#### Case 3. *fMRI*

The same linear mixed-effects modeling approach described in *Case 2* was applied to sRQA metrics from the dorsal attention network. We examined measures for subnetworks A and B, and sCRQA measures between subnetworks. Task condition (movie vs. rest) was entered as a fixed effect. Separate models were fit for 13 RQA metrics (DET, L, Lmax, DIV, ENTR, LAM, TT, Vmax, VENTR, MRT, RTE, NMPRT, TREND) and 16 cross-recurrence metrics (including additional MRT1, MRT2, RTE1, RTE2, NMPRT1, NMPRT2 metrics). FDR correction was applied across all models.

#### Case 4. Speech

The same linear mixed-effects modeling framework was applied to sRQA metrics derived from pause patterns in speech, which characterize the organization, predictability, and complexity of pauses. Statement valence (positive vs. negative) and veracity (truth vs. lie) were dummy-coded and entered as fixed effects, with participant included as a random intercept to account for repeated measures. All models controlled for sex and race as covariates. Separate models were fit for 14 sRQA metrics and FDR correction was applied to interaction-level p-values. A three-way interaction model was then estimated to examine pause patterns as a function of valence, veracity, and participant sex; lower order interactions were implicitly included and evaluated, and Tukey’s post-hoc pairwise comparisons were conducted where multiplicative effects were statistically significant.

A machine learning classification model was developed to distinguish truthful from deceptive speech using sRQA metrics. sRQA metrics were computed separately for positively- and negatively-valenced statements, and multiplicative interaction features were derived from these, resulting in 42 predictors per observation. The final dataset thus comprised 160 observations with *N* = 80 participants assessed in conditions of truthful or false speech. An XGBoost gradient boosting classifier (66,67) with DART booster was implemented, with hyperparameter optimization performed using Hyperband tuning with leave-one-out cross-validation (LOO-CV). The search space included: number of boosting rounds (10–150), learning rate (0.001-0.3), maximum tree depth (2–10), gamma (0–8), and L1/L2 regularization parameters. The final model was evaluated using LOO-CV on the complete dataset.

## DATA SOFTWARE AND AVAILABILITY

The sRQA package, along with a detailed vignette with worked examples, full function documentation, and all code used to generate the results presented in this paper (including simulation, analysis, and visualization scripts), is freely available as an open-source R library at https://github.com/lab-panda. ECG data are publicly available through the PhysioNet MIT-BIH Atrial Fibrillation Database. fMRI data are publicly available through the Reproducible Brain Chart repository. Speech data are publicly available through the Miami University Deception Detection (MU3D) dataset.

